# Spatial and mechanical environments regulate the heterogeneity of myonuclei

**DOI:** 10.1101/2024.07.08.602510

**Authors:** Rosa Nicolas, Marie-Ange Bonnin, Cédrine Blavet, Joana Esteves de Lima, Cécile Legallais, Delphine Duprez

**Affiliations:** Sorbonne Université, Institut Biologie Paris Seine, CNRS UMR7622, Developmental Biology Laboratory, Inserm U1156, F-75005 Paris, France; Univ Paris Est Creteil, Inserm, EnvA, EFS, AP-HP, IRMB, F-94010 Creteil, France; Université de Technologie de Compiègne, Sorbonne Universités, UMR CNRS 7338 Biomechanics & Bioengineering, F-60203 Compiegne, France

**Keywords:** Myonuclei, Heterogeneity, Regionalisation, Myoblast, Fibroblast, Myotendinous junction, Cell cultures, Immobilization, Limbs, Chicken, Quail, Embryos

## Abstract

Skeletal muscle formation involves tight interactions between muscle cells and associated connective tissue fibroblasts. Every muscle displays the same type of organisation, they are innervated in the middle and attached at both extremities to tendons. Myonuclei are heterogeneous along myotubes and regionalised according to these middle and tip domains. During development, as soon as myotubes are formed, myonuclei at muscle tips facing developing tendons display their own molecular program. In addition to molecular heterogeneity, a subset of tip myonuclei has a fibroblastic origin different to the classical somitic origin, highlighting a cellular heterogeneity of myonuclei in foetal myotubes. To gain insights on the functional relevance of myonucleus heterogeneity during limb development, we used 2D culture and co-culture systems to dissociate autonomous processes (occurring in 2D-cultures) from 3D-spatial environment of tissue development. We also assessed the role of mechanical parameters in myonucleus heterogeneity in paralysed limb muscles. The regionalisation of cellular heterogeneity was lost in 2D cell culture systems and paralyzed muscles. The molecular signature of MTJ myonuclei was also lost a dish and paralysed muscles indicating a requirement of spatial and mechanical input for MTJ formation. Tip genes that maintain their expression in a dish are involved in myofibrillogenesis and myotube attachments. The behaviour of regionalized markers in cultured myotubes and paralyzed muscles allows us to deduce whether the genes intervene in myogenesis, myotube attachment or MTJ formation.

**Highlights:** The regionalisation of cellular heterogeneity of myonuclei is lost in cultured myotubes

BMP signalling regulates fibroblast nucleus incorporation into cultured myotubes

The molecular signature of MTJ myonuclei is lost in cultured myotubes and paralysed muscles

Tip genes involved in myofibrillogenesis maintain their regionalised expression in cultured myotubes

## Introduction

Skeletal muscle is composed of myofibres that are multinucleated cells. Molecular and cellular heterogeneity of myonuclei has been identified in developing and adult myofibres. The functional relevance of this myonuclei heterogeneity is not fully understood for limb muscle development.

Limb muscle development involves different steps leading to a final specific limb muscle pattern. Each muscle is unique, with a specific shape, size and insertions to bone via connective tissues. The developmental programs of muscle and associated connective tissues (CT) must be tightly regulated to create a functional musculoskeletal system. In vertebrate limbs, myogenic cells and CT fibroblasts have distinct mesodermal origins. Limb myogenic cells originate from somites (paraxial mesoderm), while CT fibroblasts derive from the lateral plate mesoderm [1], [2], [3], [4]. In early limb buds, myogenic cells and fibroblasts are mixed together in dorsal and ventral limb regions. During development, myogenic cells and associated fibroblasts undergo massive spatial re-arrangements and segregate from each other to give rise to skeletal muscle and associated muscle attachments [5]. These spatial re-arrangements involve multiple molecular and cellular interactions that are not well characterized. Spatial re-arrangements are suspected to be driven by mechanical parameters, since modification of muscle contraction leads to significant defects in the musculoskeletal system formation [6], [7], [8], [9].

Although each limb muscle is unique in terms of shape and position, every muscle displays the same type of organization. They are all attached to tendons at both extremities and innervated at the centre of muscle. This defines the muscle domains, the central domain being where the innervation targets the muscle at the neuromuscular junction (NMJ) and the tip domains being at both extremities of the muscle at the myotendinous junction (MTJ). The MTJ is a peculiar structure that links two tissues with different biomechanical properties, muscle and tendon. The MTJ is mainly composed of matrix proteins produced by muscle fibres and tendon fibroblasts. Among MTJ matrix proteins, type XXII collagen is the best recognized MTJ marker at developmental, postnatal and adult stages, in mouse, zebrafish and human [10], [11], [12], [13]. *COL22A1* is mainly produced by MTJ myonuclei, type XXII collagen being deposited at the muscle/tendon interface [10], [12], [13]. In addition to being a recognized MTJ marker, *COL22A1* is required for MTJ formation and function in zebrafish [11], [14]. The molecular heterogeneity of myonuclei has been highlighted according to NMJ and MTJ domains, with the identification of NMJ and MTJ myonuclei in postnatal and adult skeletal muscles using single-nucleus RNA-sequencing strategies in mice [15], [16], [17] and human. Central NMJ myonuclei express markers of NMJ components in adult skeletal muscle, while tip MTJ myonuclei display a different molecular signature [15], [16], [17]; although with rather little overlap across studies and species [13]. In addition to molecular heterogeneity, a subset of tip myonuclei has a fibroblastic origin different to the classical somitic origin, highlighting a cellular heterogeneity of myonuclei in myofiber at the MTJ [18], [19]. Because a fibroblast genetic program has been identified in a subset of MTJ myonuclei in [16], a possible scenario would be that the fibroblasts incorporated in the myofiber at muscle tips close to tendons participate in MTJ formation.

The heterogeneity of myonuclei is observed as soon as muscle fibres are formed, with gene expression regionalized in central and tip myonuclei during development [20], [21], [6], [18], [22]. As expected, subsynaptic myonuclei at the NMJ were identified in the central domain of developing muscles [22]. Similar to NMJ genes, the fusion associated genes, *TMEM8C* (myomaker) and *MYOG* have been shown to be regionalised in the middle of limb muscles during foetal development[23]; indicative of preferential fusion events in the middle of foetal muscle close to innervation. Several signalling pathways were identified as being regionalized in tip myonuclei during development with associated function in myogenesis or MTJ formation. *FGF4* transcripts are located in tip myonuclei facing developing tendons in chicken limbs and somites [20], [24]. The secreted factor, FGF4 has been suggested to maintain the expression of tendon markers, indicative of a muscle to tendon dialogue [20], [24]. Two transcriptional readouts of YAP signalling, *ANKDR1* and *CTGF,* are regionalized in tip myonuclei in foetal muscles, with an unknown function related to muscle/tendon interface [6]. BMP signalling (assessed with pSMAD1/5/9) in tip myonuclei has been suggested to be involved in fibroblast incorporation into myotubes at the MTJ and limb muscle patterning in chicken and mouse embryos [18], [25].

The cellular and molecular heterogeneity of tip myonuclei is visualized with myonuclei of fibroblast origin and with regionalized expression of genes in developing limb muscles. Although the functional relevance of this molecular and cellular heterogeneity of myonuclei is not fully understood, it is suspected to be relevant to the different steps of the muscle program (proliferation, cell cycle exit, differentiation and fusion), myotube attachment to tendons and MTJ matrix production. In this study, we followed the molecular and cellular heterogeneity of myonuclei in cell culture systems overcoming the complexity of 3D-spatial environments and in limbs with no mechanical input (paralyzed limbs). The myoblast culture system was used to follow myonucleus heterogeneity during cell lineage progression along the myogenic program. Myoblast/fibroblast co-culture systems combined with the quail/chicken system allow the monitoring of cellular myonucleus heterogeneity i.e. fibroblast incorporation into myotubes. Embryo immobilisation allows us to assess the influence of mechanical parameters on myonucleus heterogeneity in paralyzed muscle. The comparison of cellular and molecular heterogeneity of myonuclei in a dish and paralyzed limb muscles with that of motile limb muscles allow us to deduce whether the genes intervene in myogenesis, myotube attachment or MTJ formation.

## Material and Methods

### Quail and chicken embryos

Fertilized chicken eggs (White Leghorn and JA57 strain) and quail eggs (Japanese quail strain) from commercial source (EARL Les Bruyères, Dangers, France) were incubated at 37.5°C in a humidified incubator until appropriate stages. Embryos were staged according to days of *in ovo* development. All experiments with chicken embryos were performed before E10 and consequently are not submitted to a licensing committee, in accordance with European guidelines and regulations.

### Monoculture experiments

#### Quail and chicken myoblast cultures

Primary myoblasts were isolated from limbs of E10 quail embryos or E10 chicken embryos. Limbs from quail or chicken embryos were sectioned from the main body, collected, minced and centrifuged at 1000 RPM for 1 min. Supernatant was collected and filtered through a 40 μm sieve to remove debris. Pellet was re-suspended with culture medium and centrifuged at 1000 RPM to collect the remaining cells; supernatant was collected and filtered through a 40 μm sieve. This procedure was repeated four times. Cells were distributed into culture dishes pre-coated with PBS/0,1% Porcin Gelatin. Primary myoblasts were cultured with proliferation medium (2/3 of MEM Alpha Medium, 1/3 of 199 Medium, 10% Foetal calf serum) for two days. When reaching 70% of confluence, cells were cultured in differentiation medium (MEM Alpha Medium, +1/3 199 Medium, 2% Foetal calf serum).

#### Quail and chicken fibroblast cultures

Primary fibroblasts were isolated from limbs of E10 quail embryos or E10 chicken embryos. Limbs from quail or chicken embryos were sectioned from the main body, collected and dissociated with trypsine for 20 minutes and then centrifugated for 5 minutes at 1000 RPM. The pellet was resuspended in culture media (DMEM) and plated into tissue culture dishes.

#### C2C12 cells

The mouse muscle cell line, C2C12 was cultured in a proliferation culture medium (DMEM with 4g/L glucose, 10% Foetal calf serum) to reach 50% of confluence and then in differentiation medium (DMEM with 4g/L glucose, 1% horse serum).

#### Human myoblast cell line

The human myoblast cell line AB1079 (provided by Myoline facility, Institut de Myologie, Paris) was cultured in a commercial culture medium, Skeletal Muscle Cell Growth Medium (C-23160 PROMOCELL).

#### C3H10T1/2 cells

C3H10T1/2 cells, a multipotent cell line established from mouse embryos (Reznikoff et al., 1973), was cultured in a low glucose fibroblast culture medium (DMEM with 1g/L glucose, 10% Foetal calf serum).

For monocultures, 125 000 cells were seeded per well of 6-well tissue culture plates coated with 0.1% Gelatin (approx. 13 020 cells/ cm^2^).

### Co-culture experiments with myoblasts and fibroblasts

#### Co-cultures with quail myoblasts/chicken fibroblasts or with chicken myoblasts/quail fibroblasts

For co-culture experiments, cells were seeded at a ratio of 80% myoblasts (100 000 myoblasts) and 20% fibroblasts (20 000 fibroblasts) per well. Cells were cultured in proliferation medium (50/50 of medium of each cell type) for two days and switched to differentiation medium for another three days. At day 5, cells were fixed for further analysis. <u>Co-culture experiments with human myoblasts:</u> Human myoblasts were cultured with mouse myoblasts of C2C12 cells (50% of each cell type) in 50/50 of medium of each cell type. Co-cultures with human myoblasts/C3H10T1/2 and with human myoblasts/chicken fibroblasts were performed with 80% of myoblasts and 20% of fibroblasts at Day 0 of co-culture and cultured in 50/50 of medium of each cell type for another 5 days.

### Co-culture experiments with quail myoblasts and BMP-treated chicken fibroblasts

BMP gain- and loss-of-function experiments in chicken fibroblasts were prepared as previously described [18], [21]. Briefly, primary fibroblasts from E10 chicken embryos (White Leghorn) were transfected with the following RCAS-BP(A) constructs: Empty/RCAS (control), BMP4/RCAS (containing mouse Bmp4), BMPR1Aca/RCAS (containing human BMPR1Aca, a constitutive form of BMPR1A), NOGGIN/RCAS (containing chicken NOGGIN) and BMPR1Adn/RCAS (containing human BMPR1Adn, a dominant negative form of BMPR1A) at 60% of confluence using the Calcium Phosphate Transfection Kit (Invitrogen, France). Transfected fibroblasts were cultured for 5 to 7 days to allow virus spread. The transfected fibroblasts were then mixed with quail myoblasts that were not receptive to the RCAS-BP(A) virus, with the following proportions, 80% of myoblasts and 20% of BMP-treated fibroblasts at day 0. Co-cultures were then fixed after five days of cultures and processed for in situ hybridization or/and immunohistochemistry

### Embryo immobilization

Decamethonium bromide (DMB) (Sigma) solution was freshly prepared before each experiment at 12 mM, in Hank’s solution (Sigma) with Penicillin-Streptomycin at 1% (Gibco). The control solution was prepared using Hank’s solution with 1% of Penicillin-Streptomycin. 100 µl of DMB or control solutions were administrated in chicken embryos at E7.5 and E8.5. Embryos were fixed at E8.5 (24 hours of immobilization) or E9.5 (48 hours of immobilization).

### In situ hybridization to limbs and cell cultures

Normal or immobilized embryos were fixed in 4% paraformaldehyde (Sigma-Aldrich) overnight at 4°C, then processed in 7.5%/15% gelatin/sucrose (Sigma-Aldrich) for cryostat sections. Limbs were cut in 12-μm transverse or longitudinal sections and processed for *in situ* hybridization. Alternating serial sections were hybridized with probe 1, probe 2 and probe 3 to allow comparison of expression domains on adjacent sections of the same limb.

Cell cultures were fixed in 4% paraformaldehyde (Sigma-Aldrich) overnight at 4°C. Fixed cell cultures were hybridized with one probe followed by immunohistochemistry with different antibodies.

Digoxigenin-labelled or Fluorescein-labelled mRNA probes were prepared as described for MYOD, MYOG, TMEM8C, FGF4, SCX probes [23], for ANKRD1, for MEF2C [27]. NES probe was generated from a cDNA EST probe (cambridge bioscience). COL22A1 plasmid was provided by Manual Koch, Center for Biochemistry, Cologne, Germany). Colorimetric *in situ* hybridization was performed as described [23] using an antibody against Digoxigenin labelled with alkaline phosphatase. Fluorescence *in situ* hybridization was performed as described [28], using antibodies (against Digoxigenin or Fluorescein) labelled with POD. Fluorescence was revealed with the TSA+ Cyanin 3 or 5 system (AKOYA biosciences).

### Immunohistochemistry

*In situ* hybridization to limb sections or to cells was followed with immunohistochemistry using the MF20 monoclonal antibody (DSHB cat. # MF20, undiluted supernatant), which recognized sarcomeric myosin heavy chains. Muscle progenitors were detected using the monoclonal PAX7 antibody (DSHB cat # PAX7, 1/200). The monoclonal antibodies, PAX7 and MF20 developed by D.A. Fischman and A. Kawakami, respectively were obtained from the Developmental Studies Hybridoma Bank developed under the auspices of the NICHD and maintained by the University of Iowa, Department of Biology Iowa City, IA 52242, USA. Differentiating myoblasts were detected using the MYOG antibody. The MYOG antibody was provided by Christophe Marcelle (INMG, Lyon, France). Quail nuclei were detected using the QCPN antibody (DSHB cat. # QCPN, undiluted supernatant). Active BMP signalling was detected using the polyclonal pSMAD antibody recognizing the complex BMP-activated receptor-phosphorylated SMAD1/5/9 (Cell Signalling, Ref: 9516S). Secondary antibodies were conjugated with Alexa 488, Alexa 555 or Alexa 647 (Invitrogen). Nuclei were visualized with DAPI (Sigma-Aldrich) staining.

### Image capture

After *in situ* hybridization and/or immunohistochemistry experiments images were obtained using an Apotome.2 epifluorescence microscope (Zeiss) or a Nikon eclipse E800 microscope, with the possibility to combine colorimetric and fluorescence labeling, or a Leica DMR microscope for colorimetric images only.

### Image analyses and quantification

In myoblast monocultures, the fusion index is the number of nuclei within MF20+ myotubes (containing at least 2 myonuclei) divided by the total number of DAPI+ nuclei per surface unit. Mononucleated MF20+ cells were excluded of the counting. In quail myoblast/ chicken fibroblast co-cultures, the fusion index is the number of QCPN+ nuclei within MF20+ myotubes (containing at least 2 myonuclei) divided by the total number of QCPN+ nuclei.

In quail myoblast / chicken fibroblast co-cultures, the percentage of chicken fibroblast myonuclei is the number of chicken nuclei (QCPN-negative) within MF20+ myotubes (containing at least 2 myonuclei) that were divided per the total number of myonuclei or divided per the total number of chicken nuclei (QCPN-negative). In chicken myoblast / quail fibroblast co-cultures, the percentage of quail fibroblast myonuclei is the number of QCPN+ nuclei within MF20+ myotubes (containing at least 2 myonuclei) that were divided per the total number of myonuclei or divided per the total number of QCPN+ nuclei.

The myotube area per surface unit was calculated as the MF20+ area (green area) per surface unit in quail myoblast cultures and quail myoblast / chicken fibroblast co-cultures.

In BMP gain- and loss-of-function experiments, the percentage of chicken fibroblast myonuclei is the number of chicken nuclei (QCPN-negative) within MF20+ myotubes that were divided per the total number of myonuclei in quail myoblast / chicken fibroblast (control), quail myoblast / BMPR1Aca chicken fibroblast, quail myoblast / BMP4 chicken fibroblast, quail myoblast / BMPR1Adn chicken fibroblast, quail myoblast / NOGGIN chicken fibroblast co-culture systems.

All the described quantifications were performed using the Cell counter plug-in of the free software Image J / Fiji [29].

### Statistics

GraphPad Prism 9.5.1 software was used for statistical analysis. The non-parametric test, Mann Whitney was used to determine statistical significance, which was set at p values <0.05.

## Results

To study the heterogeneity of myonuclei in different contexts, we developed 2D culture and co-culture systems and an immobilisation model in chicken embryo leading to paralyzed limb muscles.

### The fusion-associated genes lose their central location in cultured myotubes

The fusion associated genes, *TMEM8C* and *MYOG* were expressed in the middle of muscles away from tendons labelled with *SCX* expression, in chicken limb muscles (Fig. 1A-D) [23]. To monitor cell lineage progression along the myogenic program, we cultured primary quail myoblasts isolated from E10 quail limbs. After five days of culture, quail myoblasts differentiated into myotubes (Fig. 1E,F). PAX7+ and MYOG+ nuclei were present in the culture assessing the myogenic progression from PAX7+ muscle progenitors (47% of all nuclei) to MYOG+ differentiated cells (25.4% of all nuclei) (Fig. 1E-G). In five day myoblast cultures, the fusion index was of 48.7% (Fig. 1G). The differentiation gene, *MYOD* and fusion associated genes*, MYOG* and *TMEM8C* were expressed in differentiating myoblasts and myonuclei of plurinucleated myotubes of 5 day myoblast cultures (Fig. 1H-M). In contrast to limb muscles (Fig. 1A,B), *MYOG* and *TMEM8C* transcripts did not display any central location in quail and chicken cultured myotubes (Fig. 1H-M). *TMEM8C, MYOG* transcripts did not show any obvious regionalization and were enriched in zones with high density of myonuclei: at contact points between myotubes and at myotube tips (Fig. 1H,K,K’,I,L,L’).

**Figure 1.**
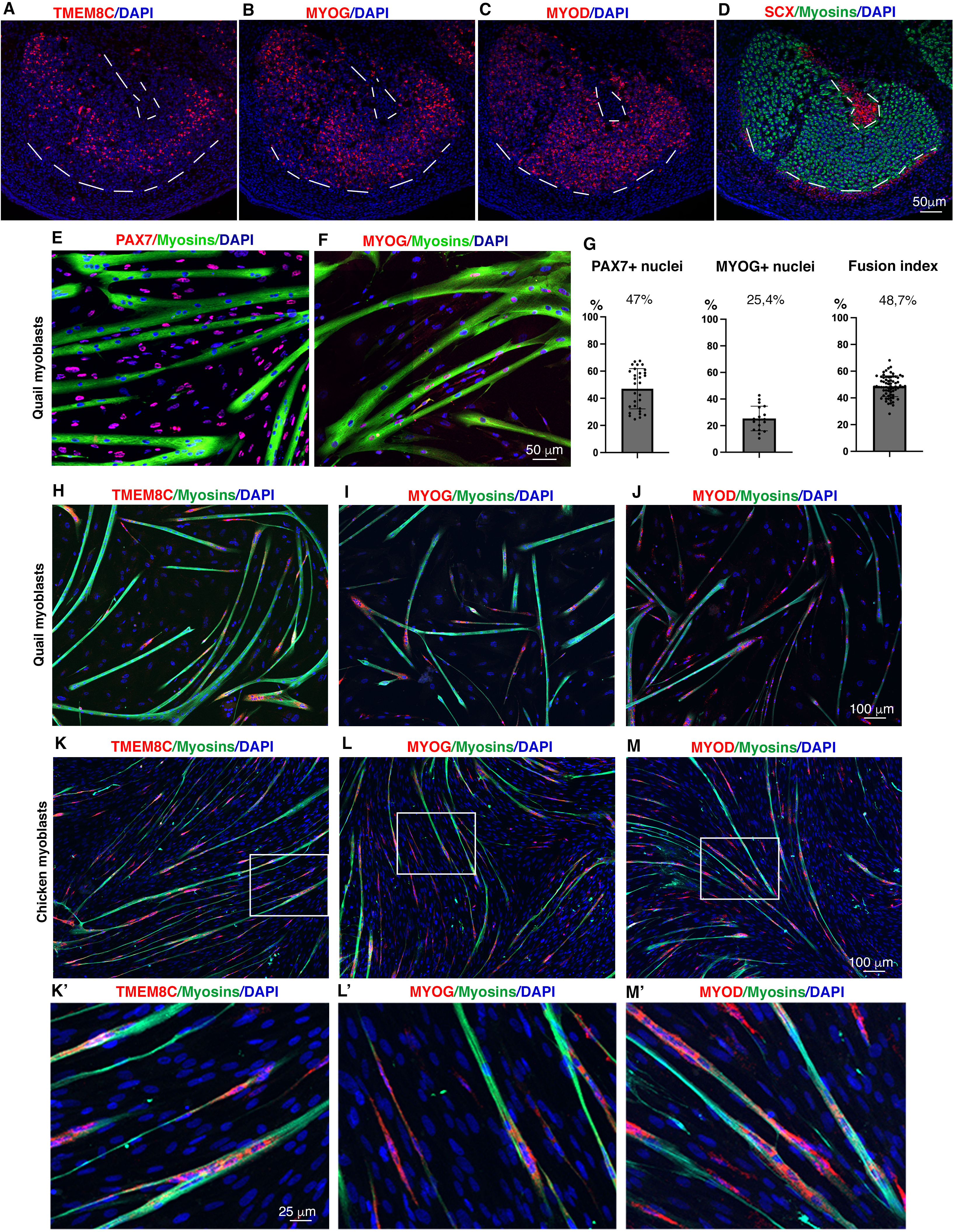
Loss of regionalisation of fusion-associated genes in cultured myotubes. (**A-D**) Fluorescent in situ hybridization to adjacent limb muscle sections from E10 chicken embryos with TMEM8C (A), MYOG (B), MYOD (C) and SCX (D) probes (red), followed by an immunolabelling of myosins (green) (D), combined with DAPI staining (nuclei, blue). (**E-J**) Primary cultures of limb myoblasts from E10 quail embryos. (**E,F**) Myoblasts and myotubes labelled with PAX7 (muscle progenitors, red) and MF20 (myosins, green) antibodies (E), or with MYOG (red) and MF20 (myosins, green) antibodies (F), combined with DAPI staining (nuclei, blue). (**G**) Percentage of PAX7-positive nuclei versus total nuclei. Percentage of MYOG-positive nuclei versus total nuclei. Fusion index. Graphs show mean±s.d. (**H-J**) Fluorescent in situ hybridization to quail myoblast cultures with TMEM8C (H), MYOG (I) and MYOD (J) probes (red) followed by an immunolabelling of myosins (green) combined with DAPI staining (nuclei, blue). (**K-M**) Fluorescent in situ hybridization to chicken myoblast cultures with TMEM8C (K), MYOG (L) and MYOD (M) probes (red) followed by an immunolabelling of myosins (green) combined with DAPI staining (nuclei, blue). (**K’-M’**) are high magnification of squared regions in (K-M).

We conclude that the fusion-associated genes lose their central location in cultured myotubes.

### The presence of fibroblasts favours all steps of the muscle program and does not change the *TMEM8C* expression pattern in cultured myotubes

To assess the effect of fibroblasts for muscle differentiation, we monitor cell lineage progression along the myogenic program differentiation in quail myoblast / chicken fibroblast co-cultures (Fig. 2). After 5 days of culture (initially plated with a ratio of 80% myoblasts and 20% fibroblasts), we observed an average ratio of 54% myoblasts and 46% fibroblasts in co-cultures (Fig. 2A); indicating that fibroblasts grow faster than myoblasts. 41.7% of PAX7+ cells and 16.5% of MYOG+ nuclei versus all nuclei (including both quail and chicken nuclei) were observed in the co-cultures (Fig. 2B-E). Because PAX7 and QCPN were both IgG1 antibodies, we could not follow PAX7+ quail nuclei. However, because there are 54% quail nuclei in co-cultures, we could estimate that the percentage of PAX7+ quail nuclei to be 77% and that of MYOG+ quail nuclei to be 30.5% versus quail nuclei (excluding chicken nuclei). So, the presence of fibroblasts induced an increase in the proportion of PAX7+ cells and MYOG+ nuclei (reported to quail nuclei) in co-cultures versus mono-cultures (47% for PAX7+ cells and 25,4% for MYOG+ nuclei, Fig. 1G). The fusion index was slightly increased in co-cultures (57.5%) versus myoblast mono-cultures (48.7%) (Fig. 2F versus Fig. 1G). We also observed an increase of myotube area (23,8%) in co-cultures compared to myoblast mono-cultures (12%) (Fig. 2G-I). The increase in the percentage of PAX7+ cells, MYOG+ nuclei, fusion index (versus quail nuclei) and myotube area/unit area in co-cultures versus mono-cultures converge to the idea that the presence of fibroblasts favours the progression of all steps of the muscle program, consistent with previous studies [30]. Consistent with the increase of MYOG+ nuclei, fusion index and myotube area/unit area, the fusion gene (Fig. 2C,E-I), *TMEM8C* is highly expressed in co-cultures (Fig. 2J,K). However, *TMEM8C* displayed the same expression pattern in myotubes of co-cultures and mono-cultures, being expressed in zones with high density of myonuclei: at contact points between myotubes and at myotube tips (Fig. 2J,K).

**Figure 2.**
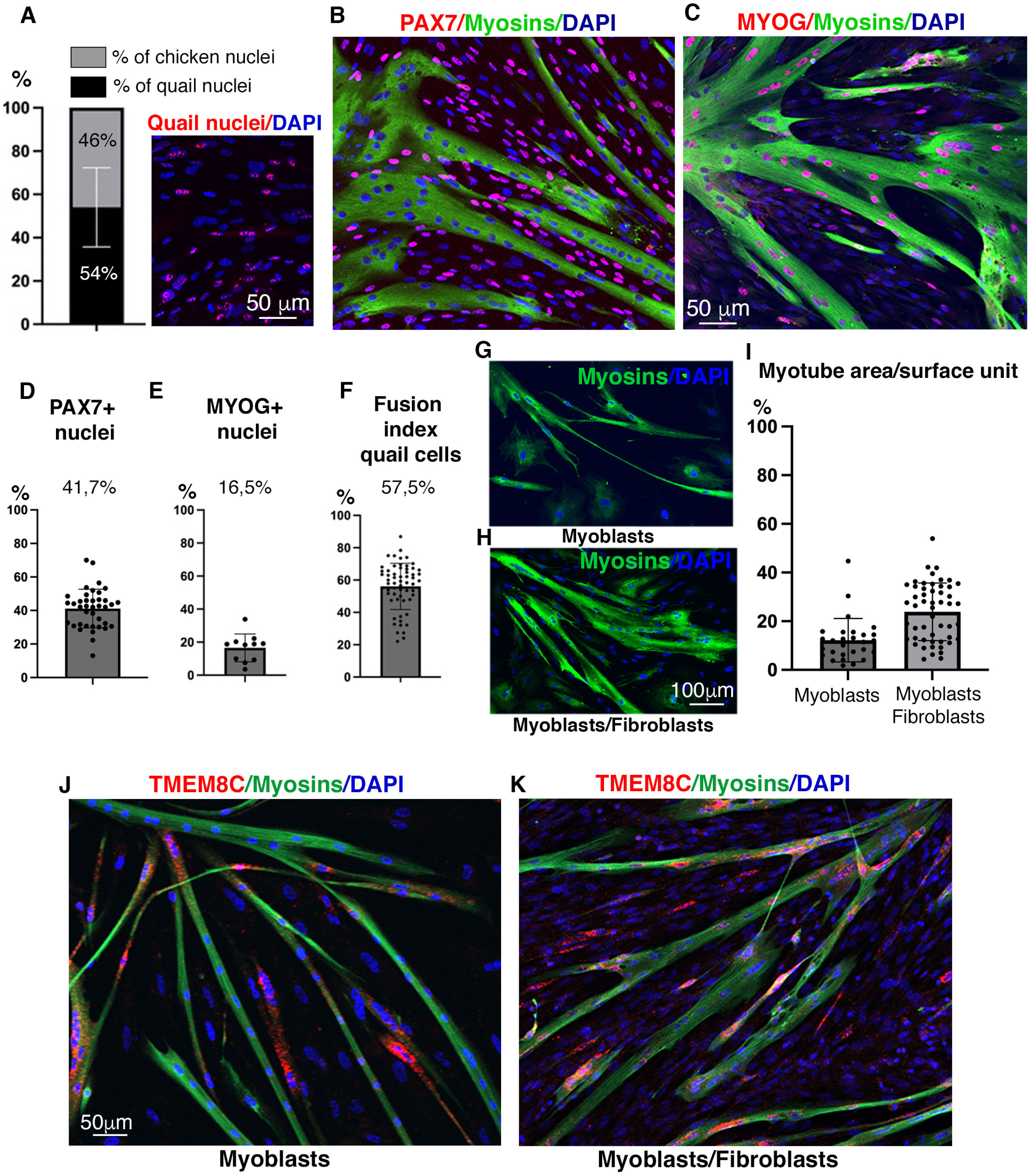
Fibroblasts promote myoblast differentiation in fibroblast/myoblast co-cultures. (**A**) Percentage of quail myoblasts and chicken fibroblasts in co-cultures after five days of cultures. Representative field of quail nuclei (QCPN antibody, red) and chicken nuclei (DAPI, blue). (**B**) Quail myoblast / chicken fibroblast co-cultures immunolabelled to PAX7 (PAX7 antibody, red) and myosins (QCPN antibody, green) combined with DAPI staining (blue). (**C**) Quail myoblast / chicken fibroblast co-cultures immunolabelled to MYOG (red) and myosins (QCPN antibody, green) combined with DAPI staining (blue). (**D,E)** Percentage of PAX7-positive nuclei (D) and of MYOG-positive nuclei (E) versus total nuclei in fibroblast/myoblast co-cultures. (**F**) Fusion index of quail cells in co-cultures. Graphs show mean±s.d. (**G,H)** Myoblast cultures (G) and myoblast/fibroblast co-cultures (H) labelled with the MF20 antibody (myosins, green), combined with DAPI staining (nuclei, blue). (**I**) Quantification of myotube area per surface unit in myoblast cultures and myoblast/fibroblast co-cultures. Graph shows mean±s.d. (**J,K**) Fluorescent in situ hybridization to quail myoblast cultures (J) and to quail myoblast / chicken fibroblast co-cultures (K) with TMEM8C probe (red) followed by an immunolabelling of myosins (green) combined with DAPI staining (nuclei, blue).

We conclude that the presence of fibroblasts promotes all steps of the muscle program and does not modify the *TMEM8C* expression pattern in myotubes of fibroblast/myoblast co-cultures.

### Fibroblast nuclei are incorporated into myotubes in myoblast/fibroblast co-cultures with no obvious regionalisation

We recently identified a cellular heterogeneity of tip myonuclei with the incorporation of fibroblast nuclei into myotubes with a preferential location at muscle/tendon interface in foetal limb muscles [18]. We first wanted to assess if fibroblast nucleus incorporation occurred in myotubes *in vitro*. To address this question, we set myoblast/fibroblast co-culture systems using the quail/chicken system that singles out quail and chicken nuclei independently to their differentiation status. We performed myoblast/fibroblast-co-culture experiments with quail primary myoblasts and chicken primary fibroblasts from E10 limbs, initially plated with a ratio of 80% myoblasts and 20% fibroblasts, and left for five days of cultures. Primary chicken fibroblasts isolated from foetal limbs did not express PAX7 or MYOG (Fig. S1A,B). Chicken fibroblasts express the SMA myofibroblast marker, the *COL12A1* connective tissue marker, and the *SCX* and *TNMD* tendon markers (Fig. S1C-F), suggestive of a fibroblast/tendon progenitor phenotype [31]. Using the QCPN antibody that recognized quail nuclei and not chicken nuclei, we followed myoblasts and fibroblasts independently of their molecular statues along the cultures. In quail myoblast / chicken fibroblast co-cultures, we appreciated a small population of myonuclei that were not of quail origin (Fig. 3A-B). Chicken fibroblast myonuclei represented 11% of the myonuclei, and 6% of fibroblast nuclei were recruited into myotubes after 5 days of culture (Fig. 3B). To exclude any potential bias regarding the labelling of quail myonuclei with the QCPN antibody, we performed the converse co-culture experiments with chicken myoblasts and quail fibroblasts. Similarly to the chicken fibroblast nucleus incorporation into quail myotubes, we observed an incorporation of quail fibroblast nuclei within chicken myotubes (Fig. 3C-C”). We quantified 20% of quail fibroblast myonuclei out of all myonuclei into myotubes and around 8% of quail fibroblast nuclei were recruited into myotubes (Fig. 3D). The fibroblast-derived myonuclei did not appear to be regionalized along the myotubes in both types of avian co-culture experiments (Fig. 3A-A”,C-C”), while fibroblast nuclei are incorporated with a preferential location at muscle/tendon interface in foetal limbs muscles [18] and postnatal muscles [19].

**Figure 3.**
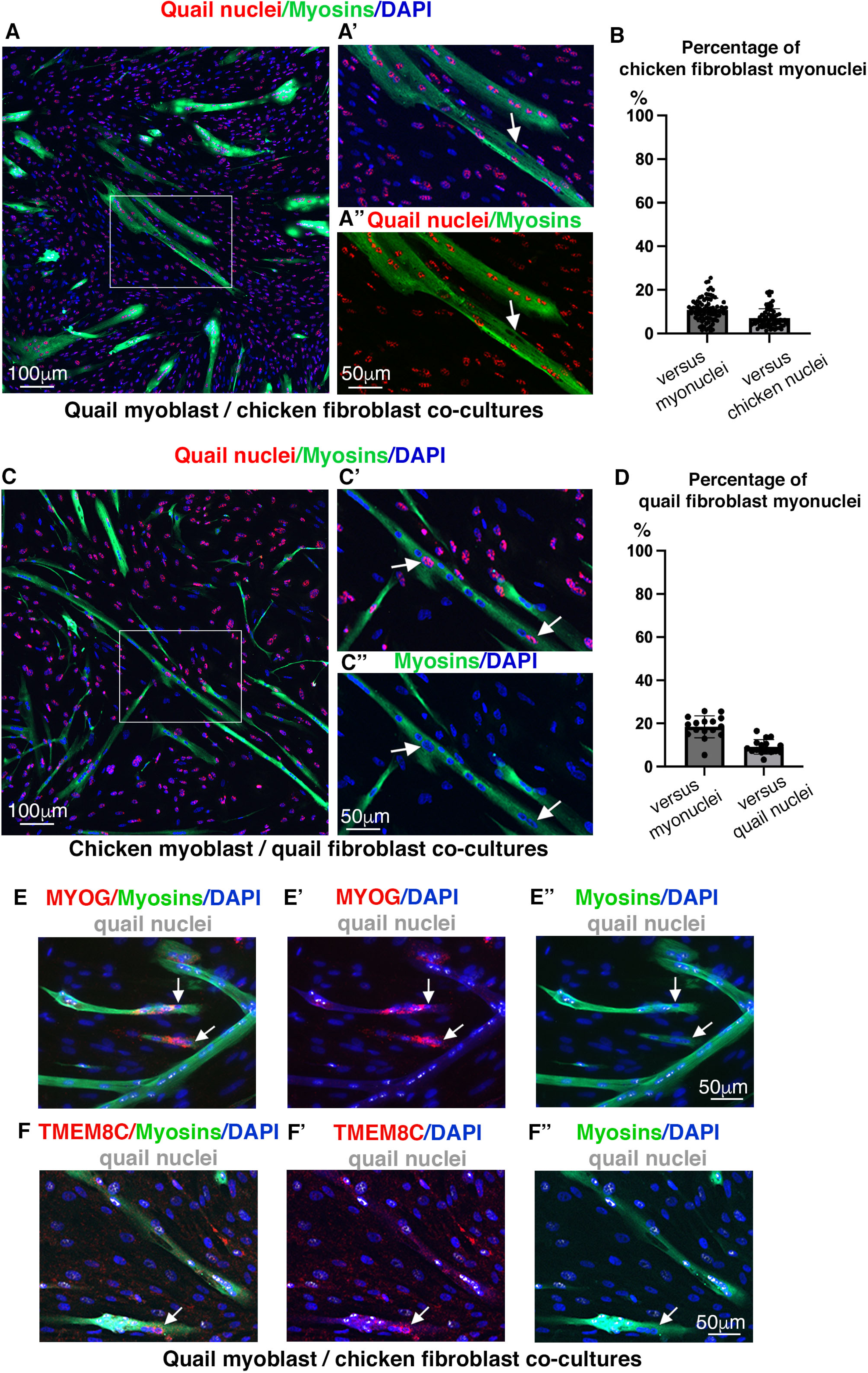
Recruitment of fibroblast nuclei into myotubes in fibroblast/myoblast co-cultures. (**A,A’,A”)** Quail myoblast and chicken fibroblast and co-cultures immunolabelled to quail nuclei (QCPN antibody, red) and myosins (MF20 antibody, green), combined with DAPI staining (nuclei, blue). (A) Left panel shows quail nuclei/Myosins/DAPI, while right panels (A’,A”) are high magnification squared in A, showing quail nuclei/Myosins/DAPI staining (A’) and quail nuclei/Myosins (A”). Arrows point to chicken fibroblast myonuclei. (**B**) Percentage of chicken fibroblast myonuclei versus quail and chicken myonuclei and versus total chicken nuclei. Graph shows mean±s.d. (**C,C’,C”**) Chicken myoblast and quail fibroblast and co-cultures immunolabelled to quail nuclei (QCPN antibody, red) and myosins (MF20 antibody, green), combined with DAPI staining (nuclei, blue). (C) Left panel shows quail nuclei/Myosins/DAPI, while right panels (C’,C”) are high magnification squared in C, showing quail nuclei/Myosins/DAPI staining (C’) and Myosins/DAPI (C”). Arrows point to quail fibroblast myonuclei. (**D**) Percentage of quail fibroblast myonuclei versus chicken and quail myonuclei and versus total quail nuclei. Graph shows mean±s.d. **(E,E’,E”,F,F’,F”**) Fluorescent in situ hybridization to quail myoblast / chicken fibroblast co-cultures with MYOG (**E,E’,E”**) or TMEM8C (**F,F’,F”**) probes (red) followed by an immunolabelling of myosins (green) and of quail cells (grey) combined with DAPI staining (nuclei, blue). Arrows point to chicken fibroblast myonuclei expressing *MYOG* (**E,E’,E”**) and *TMEM8C* (**F,F’,F”**).

In order to test if fibroblast recruitment was observed in other types of co-cultures, we performed co-cultures with a human myoblast cell line (line AB1079) (Fig. S2A-A”) associated with mouse or chicken fibroblasts. We first verified that myoblast fusion was possible between human and mouse myoblasts. Human myoblasts did fuse with mouse C2C12 myoblasts to form heterogeneous myotubes with human and mouse myonuclei (Fig. S2B-B”). Co-cultures with human myoblasts and mouse C3H10T1/2 fibroblasts (Reznikoff et al., 1973) show sporadic events of incorporation of mouse fibroblast nuclei into human myotubes (Fig. S2C-C”). We also performed co-cultures of human myoblasts with primary chicken fibroblasts and observed no or very rare events of chicken fibroblast nuclei into human myotubes (Fig. S2D-D”). We conclude that fibroblast nucleus recruitment to myotubes is observed within homologous and not heterogeneous species.

In order to assess if fibroblast myonuclei expressed the fusion-associated genes, we seek for *MYOG* and *TMEM8C* expression in fibroblast myonuclei. We found that fibroblastic myonuclei expressed the fusion-associated genes, *MYOG* and *TMEM8C* (Fig. 3E-E”,F-F”), indicating a reprograming of fibroblast myonuclei towards the myogenic program.

We conclude that fibroblast nuclei are recruited into myotubes in myoblast/fibroblast co-cultures, similarly to fibroblast recruitment into foetal limb muscles, although with no regionalisation and that the fibroblast-derived myonuclei are reprogrammed into the myogenic program.

### BMP signalling regulates the incorporation of fibroblast nuclei into myotubes in cultures

BMP signalling has been shown to regulate fibroblast nucleus incorporation into myotubes preferentially at muscle tips during limb development consistent with active BMP signalling at muscle/tendon interface [21], [18]. BMP4 ligand was produced by tendon cells (visualized with *BMP4* transcripts in tendons), while BMP-responsive nuclei, visualized with pSMAD1/5/9, were located in a subpopulation of myonuclei close to tendon, in chicken limb muscles (Fig. 4A-C). The tip regionalisation of BMP-responsive nuclei observed in limb muscles was lost in myoblast cultures since pSMAD1/5/9 was observed in all myonuclei of cultured myotubes (Fig. 4D-D”). In order to assess the effect of BMP signalling in fibroblast nucleus incorporation into myotubes, we performed BMP gain-and-loss of function experiments in chicken fibroblasts and then cultured them with quail myoblasts (Fig. 4E-J). Using the RCAS-BP(A) retroviral system, we were able to infect chicken fibroblasts and not quail myoblasts [32]. BMP gain-of-function experiments were performed with BMPR1Aca/RCAS and BMP4/RCAS [21], while BMP loss-of-function experiments were performed with BMPR1Adn/RCAS and NOGGIN/RCAS (Wang *et al.*, 2010; Nassari et al., 2017). The BMP-treated fibroblasts were then associated with quail myoblasts as co-cultures. When BMP signalling pathway was up-regulated in chicken fibroblasts, an increase of fibroblast nucleus incorporation into myotubes was observed from 11.5% to 13.3% for BMPR1Aca and from 11.5% to 14.3% for BMP4 (Fig. 4G), while the blockade of BMP signalling in fibroblasts decreased fibroblast nucleus incorporation into myotubes from 8% to 4.4% for BMPR1Adn and from 8% to 5.5% for NOGGIN (Fig. 4J). We did not observe any obvious regionalisation of fibroblast myonuclei along the myotubes in BMP treated co-cultures (Fig. 4E,F,H,I).

**Figure 4.**
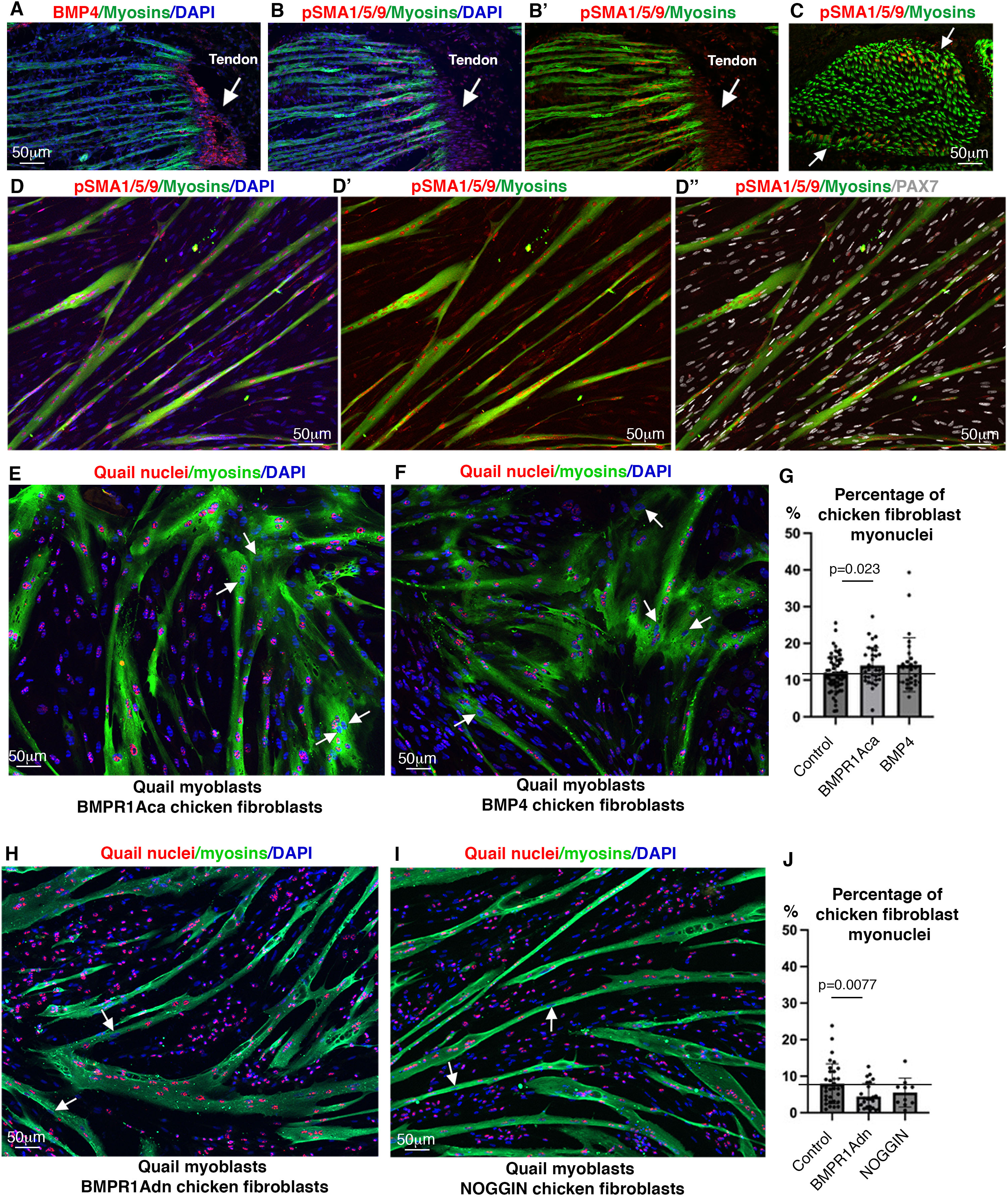
BMP signalling regulates the incorporation of fibroblast nuclei into myotubes. (**A**) Fluorescent in situ hybridization to longitudinal sections of limb muscles with BMP4 probe (red) followed by an immunohistochimistry with the MF20 antibody (myosins, green) combined with DAPI staining (nuclei, blue). (**B,B’**) Immunohistochemistry to adjacent longitudinal sections of limb muscle sections with pSMAD1/5/9 (red) and MF20 (myosins, green) antibodies combined with DAPI staining (nuclei, blue). (**C**) Immunohistochemistry to transverse sections of limb muscles with pSMAD1/5/9 (red) and MF20 (myosins, green) antibodies). (**D,D’,D”**) Immunohistochemistry to quail myoblast cultures with pSMAD1/5/9 (red), PAX7 (grey) and MF20 (myosins, green) antibodies combined with DAPI staining (nuclei, blue). (**E-J**) BMPR1Aca-transfected chicken fibroblasts (E), BMP4-transfected chicken fibroblasts (F), BMPR1Adn-transfected chicken fibroblasts (H) or NOGGIN-transfected chicken fibroblasts (I) were co-cultured with quail myoblasts; and labelled with the QCPN (quail nuclei, red), MF20 antibody (myosins, green) combined with DAPI staining (nuclei, blue). (**G**) Percentage of chicken fibroblast myonuclei within myotubes in control-, BMPR1Aca-, BMP4-transfected chicken fibroblasts cultured with quail myoblasts, (BMP gain-of-function experiments). (**J**) Percentage of chicken fibroblast myonuclei within myotubes in control-, BMPR1Adn-, NOGGIN-transfected chicken fibroblasts cultured with quail myoblasts, (BMP loss-of-function experiments). (G,J) Graphs shows mean±s.d.

We conclude that BMP signalling regulate the incorporation of fibroblasts into myotubes *in vitro*, as observed in limb muscles [18], although with no obvious regionalisation.

### Genes expressed in tip myonuclei of limb muscles behave differently in cultured myotubes

BMP-responsive myonuclei lost their tip regionalisation in cultured myotubes (Fig. 4); this prompted us to analyse in a dish, the expression of genes known to be expressed in tip myonuclei of limb muscles. We first analysed the expression of the main MTJ marker, *COL22A1* [10], [11] with that of the secreted factor, *FGF4* also expressed in tip myonuclei of developing chicken muscles [20]. Consistent with these studies, *COL22A1* and *FGF4* transcripts were located in myonuclei at muscle ends close to tendons visualized with *SCX* expression on transverse and longitudinal muscle sections of E10 chicken limbs (Fig. 5A-F). The expression of *COL22A1* and *FGF4* was lost in chicken cultured myotubes (Fig. 5G-H). No *COL22A1* expression was observed in quail cultured myotubes (Fig. 5I), while being expressed in tip myonuclei of quail limb muscles (Fig. S3). The presence of chicken fibroblasts did not induce *COL22A1* expression in quail myotubes in co-culture experiments (Fig. 5I,J).

**Figure 5.**
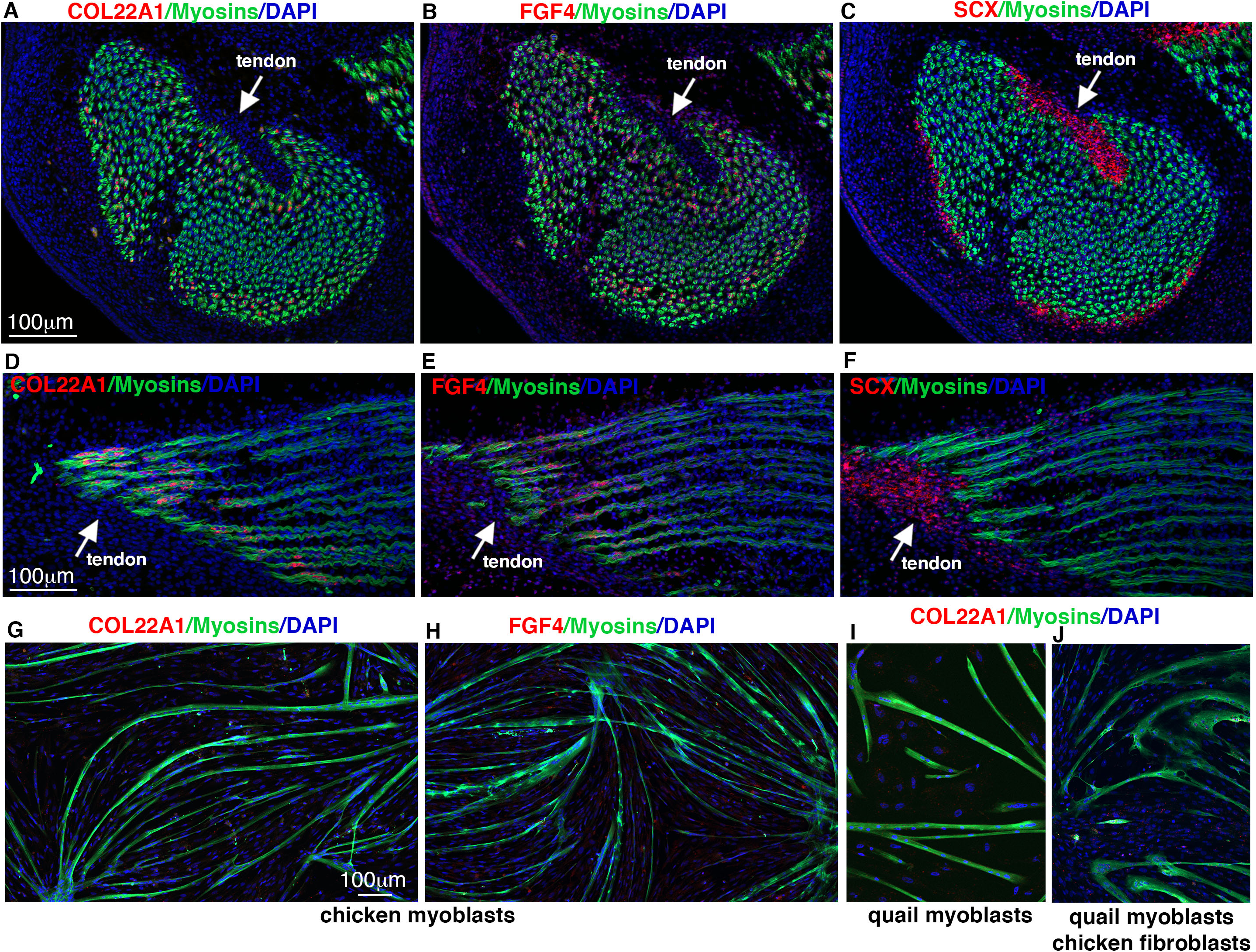
The *COL22A1* and *FGF4* expression in tip myonuclei of limb muscles is lost in cultured myotubes. (**A-F**) Gene expression for muscle tip genes (*COL22A1, FGF4*) and tendon gene (*SCX*) in adjacent transverse (A-C) and longitudinal (D-F) muscle sections from E10 chicken limbs with fluorescent in situ hybridization with COL22A1 (A,D), FGF4 (B,E) and SCX (C,F) probes (red), followed by an immunohistochemistry with the MF20 antibody to label myosins (green) combined with DAPI staining (blue). Arrows point to tendons and muscle/tendon interface. (**G-J**) Fluorescent in situ hybridization to chicken myoblast cultures (G,H), quail myoblast cultures (I) and quail myoblast / chicken fibroblast co-cultures (J) with COL22A1 (G,I,J) and FGF4 (H) probes (red) followed by an immunolabelling of myosins (green) combined with DAPI staining (nuclei, blue).

We next analysed the expression of three other genes in cultured myotubes, genes identified as being expressed in the tip domains of muscles and involved in the skeletal muscle program. *NES*, coding for the cytoskeletal intermediate filament Nestin, contributes to skeletal muscle homeostasis and regeneration in mice [33]; Nestin being located at muscle tips in limb muscles of newborn mice and adult rats [34], [35]. The Nestin protein has been shown recently to be accumulated at the myotendinous junction in human and horse muscles [36]. *ANKRD1* (ANKyrin repeat Domain 1) coding for a muscle-ankyrin repeat protein has been described as being expressed in tip myonuclei during chicken foetal development [6] and is a marker of the MTJ Col22a1+ nucleus cluster of adult mouse muscles [15]. *MEF2C* (Myocyte enhancer factor 2C) codes for a transcription factor at the crossroad of transcriptional regulations in the skeletal muscle program during muscle development, homeostasis and regeneration. *MEF2C* labels the muscle/tendon interface in axial somites during larval Xenopus development [38] and zebrafish development [39]. *NES, ANKRD1* and *MEF2C* transcripts were regionalized in tip myonuclei of foetal limb muscles of chicken embryos (Fig. 6A-C), close to *SCX* transcripts in tendon (Fig. 6D). *NES* transcripts were also regionalized in tip myonuclei of limb muscles of quail embryos (Fig. S3). In contrast to *COL22A1* and *FGF4*, the 3 tip genes, *NES, ANKRD1*, *MEF2C*, were expressed in cultured myotubes and displayed a regionalized expression pattern at myotube tips, while *MYOG* was distributed along myotubes (Fig. 6E-H). An interesting point is that these three genes were located in zones of low myosin expression, while *MYOG* transcripts were located in zones of myosin expression in cultured myotubes (Fig. 6E’-H’).

**Figure 6.**
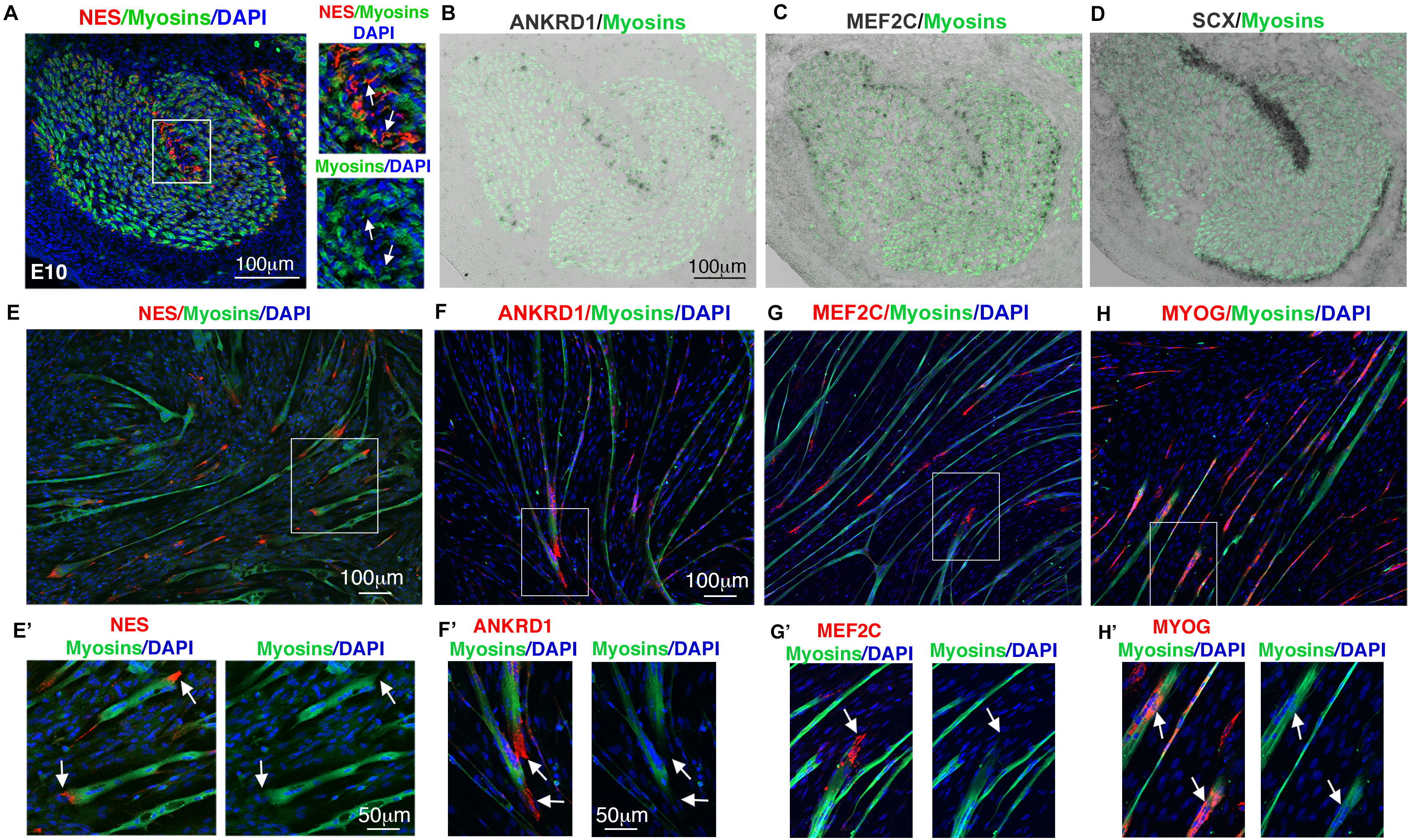
The regionalized expression of *NES*, *ANKRD1* and *MEF2C* in tip myonuclei of limb muscles is maintained in cultured myotubes. (**A-D**) Gene expression for muscle tip genes (*NES, ANKRD1, MEF2C*) and tendon gene (*SCX*) in transverse muscle sections from E10 chicken limbs with fluorescent in situ hybridization for NES probe (A) and with colorimetric in situ hybridization for ANKRD1 (B), MEF2C (C) and SCX (D) probes (dark grey), followed by an immunohistochemistry with the MF20 antibody to label myosins (green). (**E**) Fluorescent in situ hybridization to quail myoblast cultures with NES probe (red) followed by an immunolabelling of myosins (green) combined with DAPI staining (nuclei, blue). (**F-G**) Fluorescent in situ hybridization to chicken myoblast cultures with ANKRD1 (F), MEF2C (G), MYOG (H) probes (red) followed by an immunolabelling of myosins (green) combined with DAPI staining (nuclei, blue). (**E’-H’**) are high magnifications of squared areas in (E-H). Arrows point to *NES* (E’), *ANKRD1* (F’) and *MEF2C* (G’) expression at myotube tips in low myosins-expressing zones, while *MYOG* transcripts (F’) are expressed in myosins-positive zones.

In order to assess if the presence of fibroblasts affects the regionalisation of tip genes in myotubes, we analyse the expression of *NES, ANKRD1* and *MEF2C* in co-culture experiments with quail myoblasts and chicken fibroblasts. The presence of fibroblasts did not modify the tip expression of *NES, ANKRD1* and *MEF2C* transcripts in cultured myotubes (Fig. 7A-C). Thanks to the quail/chicken system, we could assess *NES* expression in fibroblast-derived myonuclei. The fibroblast-derived myonuclei did or did not express *NES*, consistent with the absence of regionalization of fibroblast recruitment and the persistent *NES* expression in myotube tips of co-cultures. (Fig. 7D,D’,E, F-F”).

**Figure 7.**
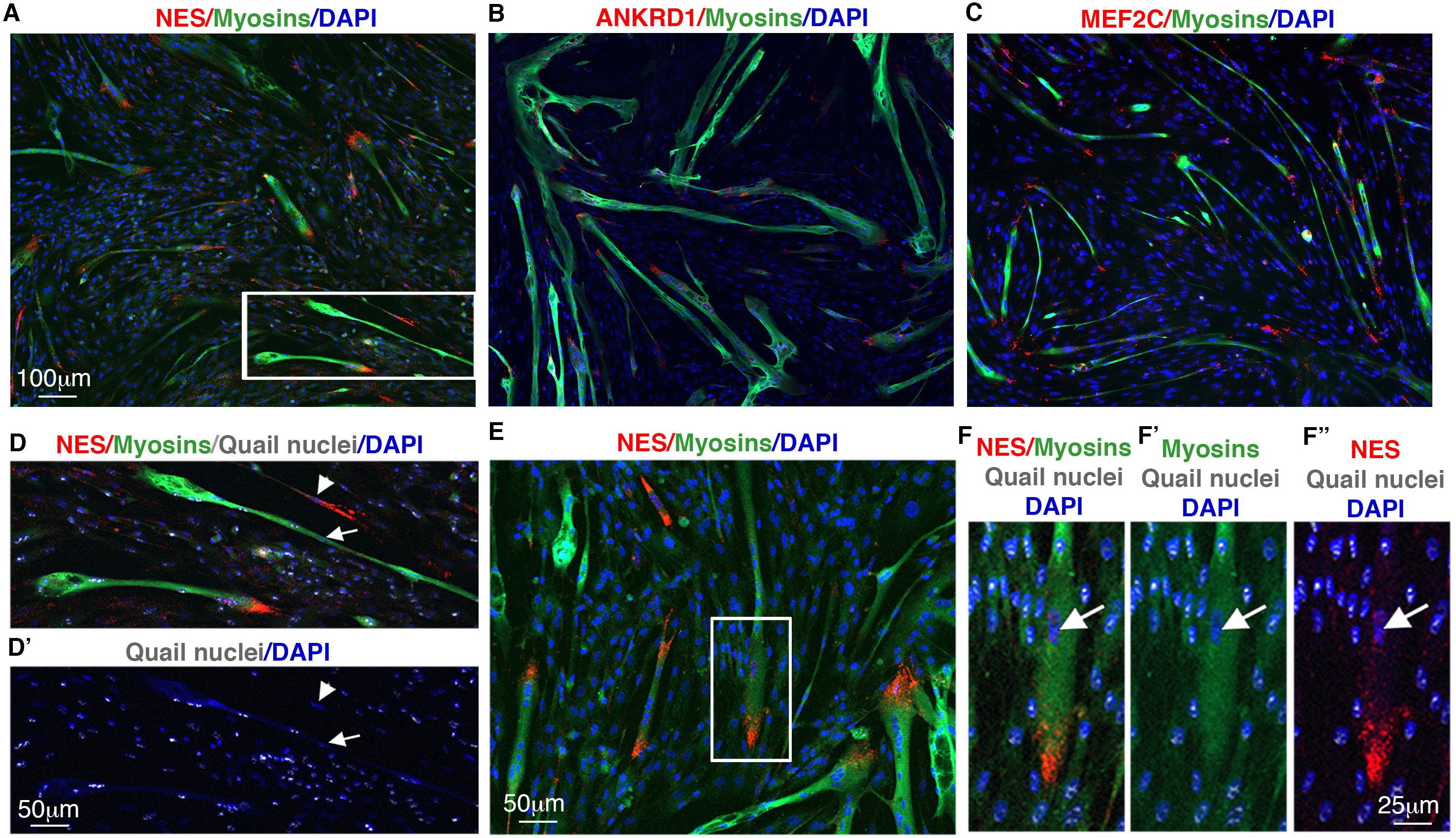
The regionalized expression of *NES*, *ANKRD1* and *MEF2C* in cultured myotubes is not changed in the presence of fibroblasts. (**A-C**) Fluorescent in situ hybridization to quail myoblast / chicken fibroblast co-cultures with the NES (A), ANKRD1 (B) and MEF2C (C) probes (red), followed by an immunohistochemistry with the MF20 antibody to label myosins (green) combined with DAPI staining (blue). (**D,D’)** High magnification of the area squared in A, showing in the top panel, (NES transcripts, red, myosins, green) and DAPI staining, blue) combined with immunolabelling of quail nuclei (grey) (D), and in the bottom panel, immunolabelling of quail nuclei (grey) combined with DAPI staining, (blue) (D’). Arrowheads point to chicken fibroblast myonuclei expressing *NES*, while arrows point to point to chicken fibroblast myonuclei not expressing *NES*. **(E**) Fluorescent in situ hybridization to quail myoblast / chicken fibroblast co-cultures with NES probes (red) followed by an immunolabelling of myosins (green) and of quail nuclei (grey), combined with DAPI staining (nuclei, blue). (**F,F’,F”**) High magnification of the area squared in E, showing NES transcripts, (red), myosins, (green) and DAPI staining (blue) combined with immunolabelling of quail nuclei (grey) (F), showing myosins, (green) and DAPI staining (blue) combined with immunolabelling of quail nuclei (grey) (F’) and showing NES transcripts, (red), DAPI staining (blue) combined with immunolabelling of quail nuclei (grey) (F”). (F,F’,F”) Arrows points to a chicken fibroblast myonuclei with low/residual levels of *NES* incorporated into a quail myotube.

We conclude that the tip markers display three types of behaviours in cultured myotubes: loss of regionalisation (pSMAD1/5/9), loss of expression (*COL22A1*, *FGF4*), or maintenance of regionalized expression (*NES, ANKRD1*, *MEF2C*). The presence of fibroblasts does not modify the expression pattern of tip markers in cultured myotubes.

### Inhibition of muscle contraction affects gene expression in tip myonuclei of limb muscles

Mechanical parameters are recognized to be an important regulator of the musculoskeletal system[8]. Muscle contractions have been shown to regulate the steps of the muscle program during foetal myogenesis [6], [23]. The absence of muscle contraction reduces the pool of muscle progenitors, while increasing their propensity to differentiate and fuse during foetal myogenesis. In order to assess the behaviour of genes expressed in tip myonuclei in absence of mechanical signals, we analysed the expression of tip genes in an unloading model during foetal myogenesis. We used the decamethonium bromide (DMB), which blocks muscle contractions and leads to limb muscle paralysis. Two days after the inhibition of muscle contraction, the expression of *COL22A1* and *FGF4* was lost in limb muscles (Fig. 8A-D), reminiscent of the loss of *COL22A1* and *FGF4* expression in myotube cultures (Fig. 5G-I). The regionalisation of pSMAD1/5/9 in tip myonuclei in foetal limbs was lost in limbs immobilized embryos (Fig. S4); reminiscent of pSMAD1/5/9 location in all myonuclei of cultured myotubes (Fig. 4D-D’’). Lastly, the regionalised expression of *NES* was maintained in limb muscles in the absence of muscle contraction (Fig. 8E,E’,F,F’), reminiscent of the regionalized expression of *NES* in cultured myotubes (Fig. 6E,E’), while the expression of *ANKRD1* [6] and *MEF2C* (Figure S5) was lost in limbs immobilized embryos.

**Figure 8.**
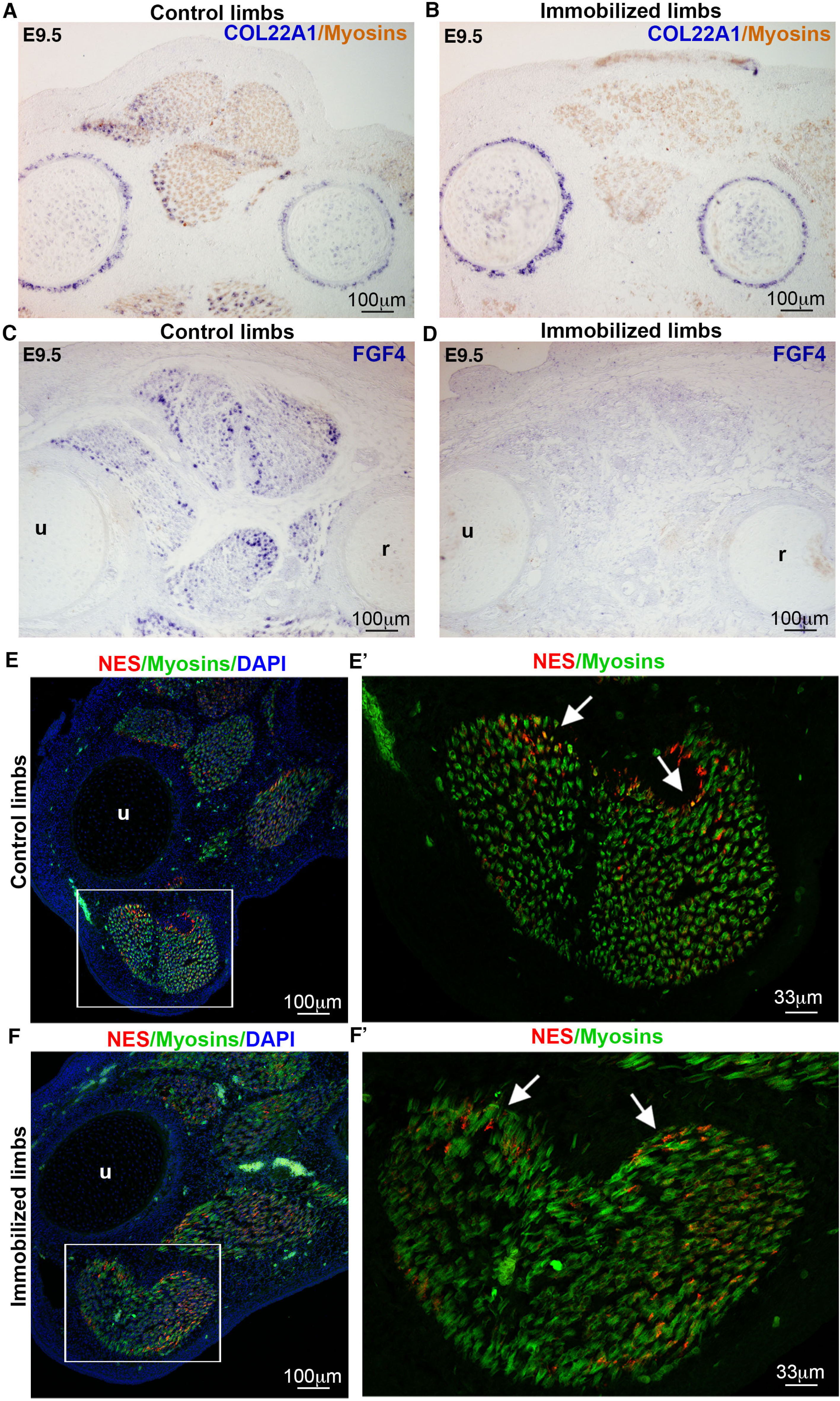
Inhibition of muscle contraction modified the expression of genes expressed in tip myonuclei of foetal muscles. Chicken embryos were treated with DMB at E7.5 to block muscle contraction and analysed 48 hours after treatment at E9.5 (N=3). (**A-D**) In situ hybridization to transverse limbs sections at the level of the zeugopod of control (A,C) and DMB-treated (B,D) E9.5 chicken embryos, with COL22A1 (A,B) and FGF4 (C,D) probes (blue) followed by immunostaining with the MF20 antibody to label myosins (light brown). The expression of *COL22A1* and *FGF4* was lost in paralyzed muscles (B,D) compared to control muscles (A,C). Note that *COL22A1* expression is maintained in cartilage and perichondrium in paralyzed limbs (B) as in control limbs (A). (**E, E’, F,F’**) Fluorescent situ hybridization to transverse limbs sections at the level of the zeugopod of control (E,E’) or DMB-treated (F,F’) E9.5 chicken embryos, with NES probe (red) followed by immunostaining with the MF20 antibody to label myosins (green) and stained with DAPI (blue). (E’,F’) are high magnification of the FCU muscles of E (control) and F (DMB-treated) limbs. u, ulna, r, radius.

We conclude that the mechanical signals affect the molecular heterogeneity of myonuclei in limb foetal muscles.

## Discussion

Myonucleus heterogeneity is altered in a dish and paralyzed limb muscles versus (motile) limb muscles. This shows the requirement of spatial and mechanical environments for myonucleus heterogeneity. The different behaviours of regionalised markers in the absence of spatial and mechanical environments also allow us to deduce whether the genes intervene in myogenesis, myotube attachment or MTJ formation.

### The regionalisation of cellular heterogeneity of myonuclei is dependent of spatial and mechanical constraints

Cellular heterogeneity of myonuclei is assessed with the regionalization of myoblast fusion in the middle and fibroblast incorporation at the tips during muscle development.

The regionalized expression of the fusion associated genes, *TMEM8C* and *MYOG* in limb muscles is indicative of a preferential location of myoblast fusion in central domains of limb foetal muscles [23]. The central regionalisation of *TMEM8C* and *MYOG* transcripts along myotubes was lost in cultures (Fig.1H,I,K,L). This indicates that the regionalisation of *TMEM8C*-dependent myoblast fusion is dependent of a spatial environment not present in a dish. The presence of fibroblasts in co-culture experiments did not change the loss of regionalization of the fusion genes along myotubes, but increased myoblast fusion (Fig. 2J,K). Interestingly, the absence of mechanical stimuli also leads to an increase of myoblast fusion and to a loss of *TMEM8C* and *MYOG* regionalized expression in paralyzed limb muscles (Esteves de Lima et al., 2016, 2022). It was recently shown that TGFβ inhibition promotes myoblast fusion *in vivo* and *in vitro* [40]. Although TGFβ ligands are expressed in limb muscles with no clear specific regionalisation [7], we cannot exclude a regionalization of TGFβ activity within muscles using the heterodimer receptor complementation mechanism [40].

In addition to be supportive actors of myogenesis, fibroblasts contribute to myotube formation. A contribution of fibroblast nuclei to myotubes has been recently highlighted during myogenesis during foetal development and postnatal life [18], [19], [41], [42]. We recapitulated the incorporation of fibroblast nuclei into myotubes in avian co-culture experiments; consistent with the incorporation of Prrx1 lineage into myotubes in mouse co-culture experiments [19]. This indicates that fibroblast incorporation into myotubes is a robust phenomenon independent of any spatial environment. Fibroblast nuclei incorporate myotubes at a ratio of about 12% of all myonuclei in avian co-culture experiments. This is very similar to the ratio observed in developing chicken and mouse limb muscles [18]. The fact that fibroblast myonuclei expressed the *TMEM8C* and *MYOG* genes (Fig. 3E,F) suggests that fibroblast myonuclei are reprogrammed into the myogenic program. In contrast to limb muscles, the incorporation of fibroblast nuclei was not regionalized in cultured myotubes and BMP signalling was active in all myonuclei of cultured myotubes. As in foetal limb muscles, BMP signalling regulates the incorporation of fibroblast nuclei in co-culture experiments (Fig. 4). There is a striking correlation between active BMP signalling and fibroblast incorporation, with no regionalisation along myotubes in culture experiments and regionalisation at the tips of myotubes within foetal muscles, strengthening BMP function in the incorporation of nuclei of fibroblast origin into myotubes *in vitro* and *in vivo*. The regionalisation of BMP signalling was lost in paralyzed muscles, indicating a requirement of mechanical input for fibroblast incorporation in foetal libms.

We conclude that the regionalisation of cellular heterogeneity of myonuclei is lost in 2D-culture systems and paralysed limb muscles.

### MTJ myonuclei lose their molecular signature in 2D-culture systems and immobilisation conditions

*COL22A1* the recognized marker of MTJ myonuclei, involved in MTJ formation and homeostasis [11], [14] is a powerful proxy for the MTJ. *COL22A1* expression was not observed in cultured myotubes and not induced in myonuclei of myoblast/fibroblast co-cultures, indicative of an absence of MTJ in 2D-culture systems. *COL22A1* was also lost in tip myonuclei of foetal limb muscles of immobilized embryos. The similar loss of *FGF4* expression in cultured myotubes and paralyzed muscles is consistent with the identified function for FGF4 to regulate the expression of tendon collagen genes downstream of mechanical forces [7], [20]. Based on *COL22A1* and *FGF4* behaviour in paralyzed limbs and culture systems, we conclude that spatial environment and mechanical input are required to generate a muscle/tendon interface at the origin of MTJ. This is consistent with the requirement of mechanical forces for MTJ formation in mice [43]. We speculate that a 2D- or 3D-environment with controlled mechanical forces will be more suitable to generate a muscle/tendon interface that resemble to MTJ.

### Genes that maintain expression in tip myonuclei in cultured myotubes are involved in myofibril formation

In addition to *COL22A1*, which has a recognized function in MTJ formation, we identified other genes expressed in tip myonuclei during foetal myogenesis. We identified three tip markers (*MEF2C, ANKRD1, NES*) of limb foetal muscles, which kept their regionalized tip expression in cultured myotubes. In contrast to the *COL22A1* MTJ proxi, the expression of these genes is independent of spatial environment. The proteins coded by these three genes are *a priori* unrelated in term of structure, one transcription factor (MEF2C), one thick-filament-binding protein shuttling from cytoplasm to nucleus (ANKRD1), and one intermediate filament (Nestin). However, these three proteins have in common being linked to myofibril formation in muscle fibres. MEF2C has been shown to be required for the assembly of myosin-containing thick filaments in nascent muscle fibres [44]. ANKRD1 acts as a junctional adhesion protein maintaining the integrity of striated muscle sarcomeres through its connection with two myofilaments: titin and myopalladin ([45], [46], [47]. Lastly, Nestin location is modified in a defective myofibril assembly mouse model [35]. Interestingly, the enzyme LoxL3 expressed at muscle tips and cultured myotubes is involved in the anchoring of myotubes to tendon matrix at the MTJ in mice [48]. We speculate that the tip genes that maintain their expression in a dish could be involved in the process of attachment of sarcomeres at myotube tips to the culture dish. Because culture plates have been plated with gelatin (a denatured collagen), it is tempting to speculate that during development these tips genes are the bridging link in the attachment of sarcomeres at myotube tips to matrix components produced by tendon fibroblasts in limb muscles.

In addition to being linked to myofibrillogenesis of skeletal muscles, ANKRD1and MEF2C are also linked to mechanical stress inherent to sarcomere contractile function. ANKRD1 functions as a mechano-sensing protein along the titin protein for skeletal muscle remodelling [49] and regulates passive forces by locking titin to thin filaments, protecting sarcomere from mechanical damage [50]. MEF2C translation has been shown to be controlled by muscle activity to regulate myofibrillogenesis [39]. Consistent with this mechano-sensing function, the *ANKRD1* and *MEF2C* expression is lost in paralysed muscles of immobilized embryos (Esteves de Lima *et al.*, 2016, Fig. S5). The maintenance of *ANKRD1* and *MEF2C* expression in tips of cultured myotubes can be explained by their additional functions in physical attachment of myotubes to the culture dish. In contrast to *ANKRD1* and *MEF2C*, the regionalized expression of *NES* was still observed in paralyzed muscles, suggesting an expression independent of the loss of mechanical signals during foetal myogenesis. However, the *NES* mRNA levels were increased at the MTJ after exercise, while those of *COL22A1* were unaffected in human mature MTJ [51]. The *NES* mRNA level increase after exercise is to be related to the increase of sarcolema length at the MTJ in rat muscles in trained conditions [52], [53].

## Conclusion

In the absence of spatial and mechanical constraints, we observed different types of behaviour for the regionalized markers of myonuclei. These different behaviours provide us with clues to attribute functions to regionalized markers, in myogenesis, myotube attachment and/or MTJ formation. A first behaviour was the loss of regionalisation of myonucleus markers in a dish and paralysed limbs. This concerns the fusion-associated genes, *TMEM8C*, *MYOG*, and the regulator of fibroblast incorporation, pSMAD1/5/8. Because these regionalized markers are linked to cellular heterogeneity, myoblast fusion in the middle domain and fibroblast incorporation in the tip domains, we conclude that the regionalization of cellular heterogeneity of myonuclei is dependent on mechanical and spatial constraints. A second behaviour was a complete loss of gene expression in cultured myotubes and paralyzed muscles for the tip genes, *COL22A1* and *FGF4*. We believe that the tip genes that lose expression in the absence of spatial constraints and mechanical input are involved in MTJ formation. A third behaviour was the maintenance of regionalized expression in tip myonuclei of cultured myotubes with the intermediary filament nestin (*NES)*, the thick-filament-binding protein, ANKRD1 and the transcription factor MEF2C. The common denominators of these tip genes converge to a function in myofibrillogenesis and in the anchoring of myotubes to matrix.

These results highlight the importance of spatial and mechanical environments for the molecular and cellular heterogeneity of myonuclei and shed light about the relevance of this heterogeneity for myogenesis, myotube attachment and/or MTJ formation.

## Aknowledgement

We thank Estelle Hirsinger, Marie-Claire Delfini and Bruno Della Gaspera for critical reading the manuscript.

## Author contribution

DD conceived the project. RN, CB, MAB, JEdL performed the experiments. RN, JEdL, DD analyzed the data. RN wrote the original draft. DD, RN, JEdL, CL reviewed and edited the original draft. DD and CL supervised the project and acquired funding.

