## Supplementary material for "Spatial and mechanical environments regulate the heterogeneity of myonuclei": Figures_Sup

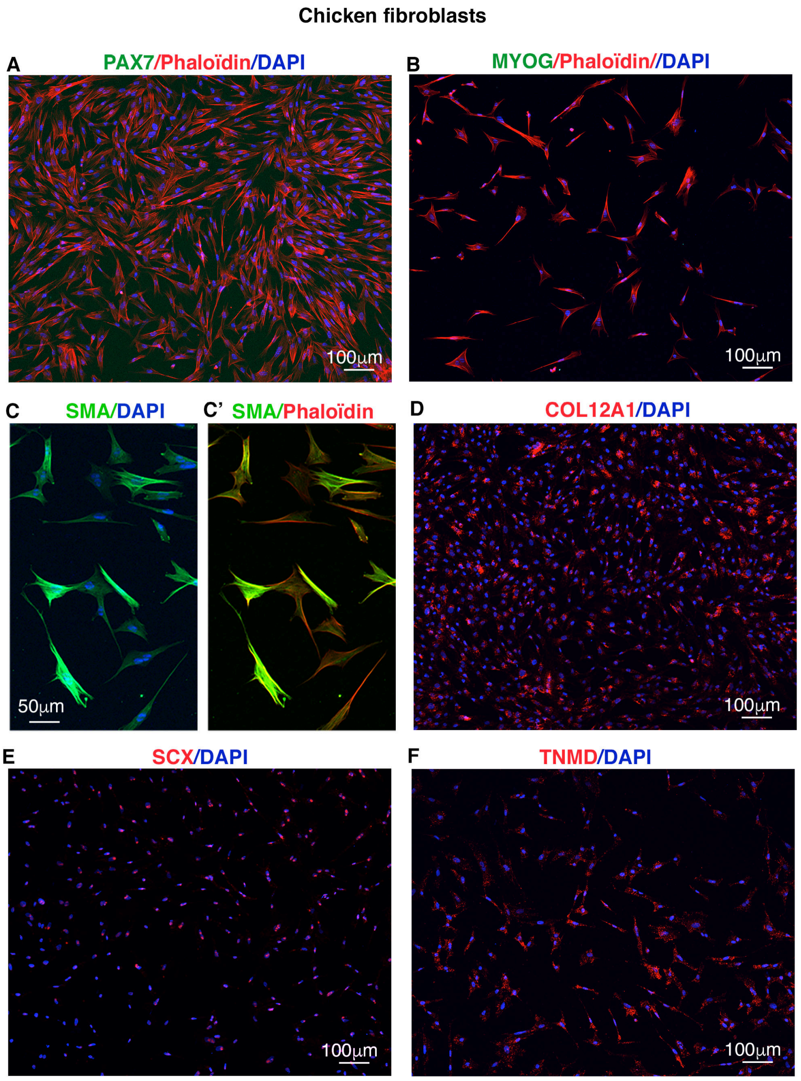


**Figure S1**

**Primary cultures of limb fibroblasts from E10 chicken embryos.**
(**A,B**) Representative fields of fibroblasts labelled with PAX7 antibody (green, A) or with MYOG antibody (green, B) combined with Phalloidin (red) and DAPI (Blue) staining. (**C,C’**) Representative fields of fibroblasts labelled with SMA antibody (green) combined with DAPI (Blue) staining (C), or with SMA antibody (green) combined with Phalloidin (red) (C’). **(D,E,F)** Representative fields of fibroblasts labelled with fluorescent in situ hybridization to COL12A1 (D), SCX (E) or TNMD (F) probes (red). (D-F) Note the nuclear location of *SCX* transcripts (E) and the cytoplasmic location of *COL12A1* (D) and *TNMD* (F) transcripts in fibroblasts.

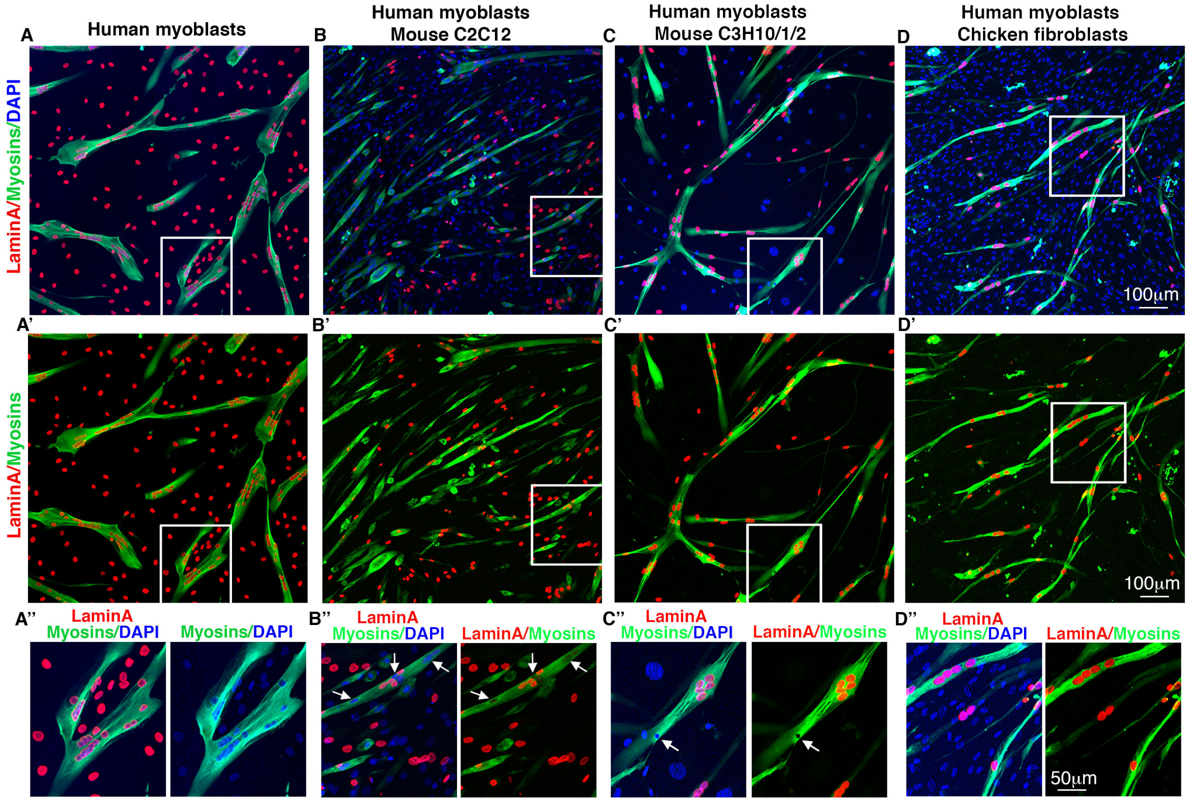


**Figure S2**

**Human myoblast / fibroblast co-cultures**

**(A,A',A")** Cultures of human myoblasts immunolabelled with the LaminA antibody (red) and the MF20 antibody to label myosins, combined with DAPI staining to label nuclei (blue). Panel A shows LaminA (red), myosin (green) and DAPI (blue) staining. Panel A' shows LaminA (red) and myosin (green) staining. Panel A" is a high magnification of squared in (A,A').

**(B,B',B")** Co-cultures of human myoblasts and mouse C2C12 myoblasts immunolabelled with the LaminA antibody (red) and the MF20 antibody to label myosins, combined with DAPI staining to label nuclei (blue). Panel B shows LaminA (red), myosin (green) and DAPI (blue) staining. Panel B' shows LaminA (red) and myosin (green) staining. Panel B" is a high magnification squared in (B,B'). Arrows point to mouse C12C12 myonuclei, LaminA negative myonuclei (blue).

**(C,C',C")** Co-cultures of human myoblasts and mouse C3H10T/1/2 fibroblasts immunolabelled with the LaminA antibody (red) and the MF20 antibody to label myosins, combined with DAPI staining to label nuclei (blue). Panel C shows LaminA (red), myosin (green) and DAPI (blue) staining. Panel C' shows LaminA (red) and myosin (green) staining. Panel C" is a high magnification squared in (C,C'). Arrows point to a fibroblast myonucleus, LaminA negative myonuclei (blue).

**(D,D',D")** Co-cultures of human myoblasts and chicken fibroblasts immunolabelled with the LaminA antibody (red) and the MF20 antibody to label myosins, combined with DAPI staining to label nuclei (blue). Panel D shows LaminA (red), myosin (green) and DAPI (blue) staining. Panel D' shows LaminA (red) and myosin (green) staining. Panel D" is a high magnification squared in (D,D').

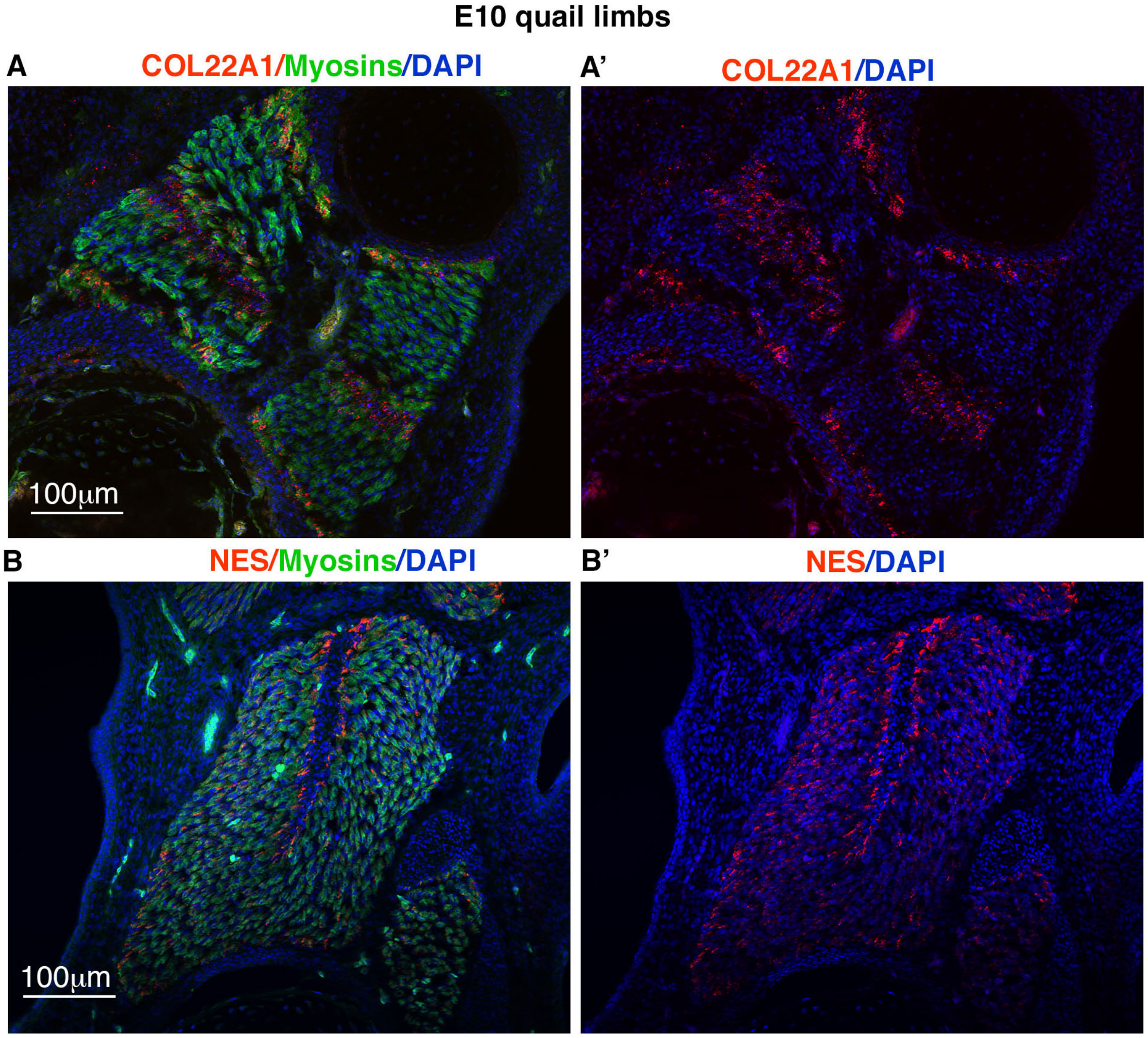


**Figure S3**

**Tip gene expression in quail limbs**
(**A,A'**) Fluorescent in situ hybridization to transverse limb sections of E10 quail embryos with the COL22A1 probe (red), followed by an immunohistochemistry with the MF20 antibody to label myosins (green), combined with DAPI staining (blue). A' is the same photograph as A and shows the *COL22A1* and DAPI staining.
**(B,B')** Fluorescent in situ hybridization to transverse limb sections of E10 quail embryos with the NES probe (red), followed by an immunohistochemistry with the MF20 antibody to label myosins (green), combined with DAPI staining (blue). B' is the same photograph as B and shows the *NES* and DAPI staining.

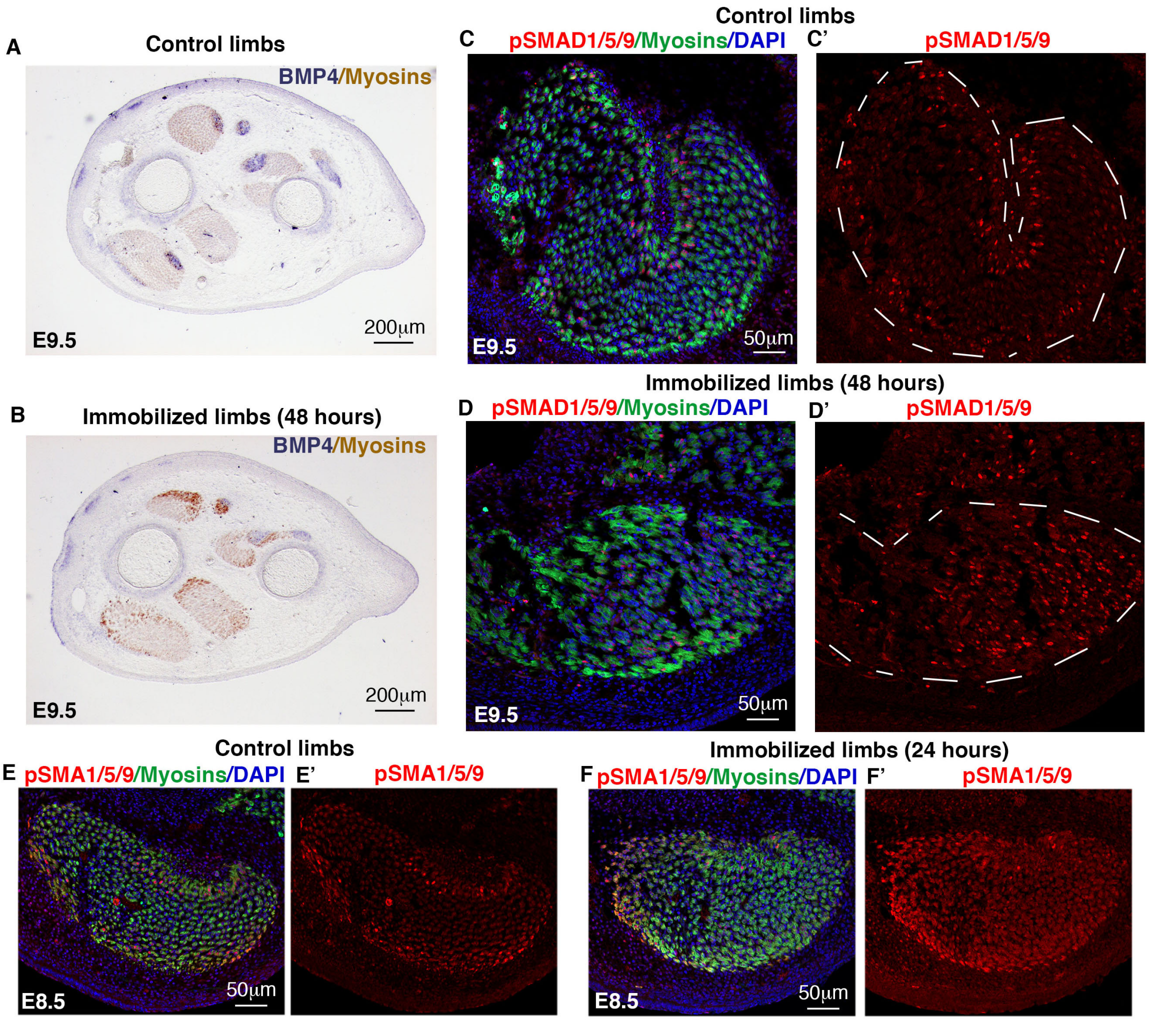


### Figure S4

**BMP signalling in limbs in the absence of muscle contraction**

**(A,B)** In situ hybridization to transverse limbs sections with BMP4 (blue) followed by immunostaining with the MF20 antibody to label myosins (light brown) for control (A, N=3) and DMB-treated limbs treated for 48 hours (B, N=3). BMP4-tendon expression is downregulated in paralyzed muscles (A,B). **(C,C',D,D')** Transverse muscle sections were immunostained with the pSMAD1/5/9 antibody (red), the MF20 antibody to label myosins (green) and stained with DAPI to visualized nuclei (blue) in control (N=3, C,C') and DMB-treated limbs for 48 hours (N=3, D,D'). White dashed lines delineate muscles. **(E,E',F,F')** Transverse muscle sections were immunostained with the pSMAD1/5/9 antibody (red), the MF20 antibody to label myosins (green) and stained with DAPI to visualized nuclei (blue) in control (N=3, E,E') and DMB-treated limbs for 24 hours (N=3, F,F').

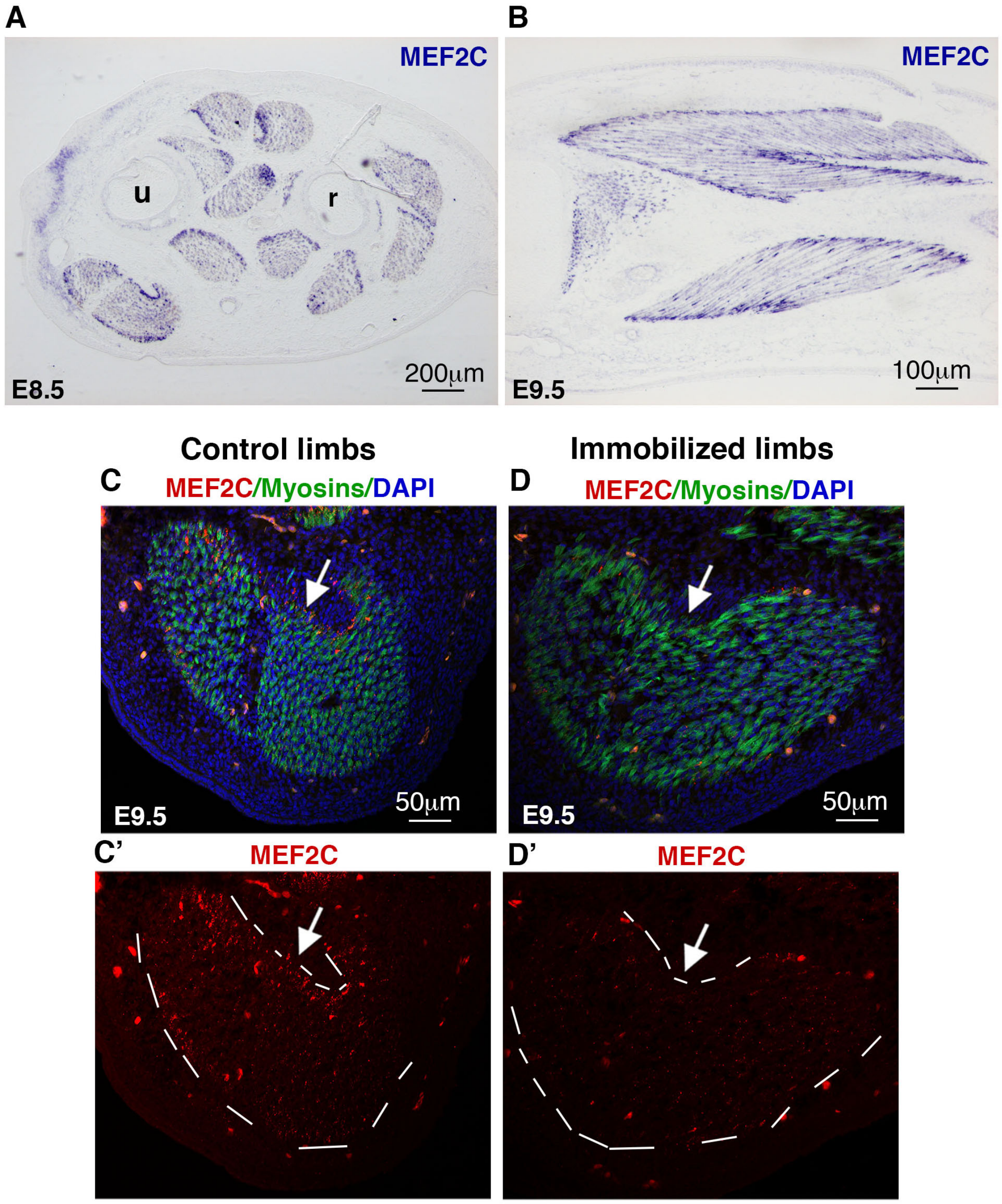


**Figure S5**

**MEF2C expression in the absence of muscle contraction**

(**A,B**) Colorimetric in situ hybridization to transverse (A) and longitudinal (B) limb sections with MEF2C probe (blue) followed with immunostaining with the MF20 antibody (light brown) of E8.5 and E9.5 chicken embryos. (**C,C',D,D'**) Fluorescent in situ hybridization to transverse muscle sections with MEF2C (red) followed by immunostaining with the MF20 antibody to label myosins (green) and stained for DAPI (blue) for control limbs (N=3, C,C') and DMB-treated limbs for 48 hours (N=3, D,D'). (C,D) show the MEF2C/Myosins/DAPI staining. (C',D') show the same sections (C,D) with MEF2C only-staining. (C',D') The white dashed lines delineate muscle shapes. u ulna, r, radius. (C,C',D,D') Arrows point to tip *MEF2C* expression in control muscles (C,C') and to loss of MEF2C expression (D,D') in paralyzed muscles.
